# Aging and injury drive neuronal senescence in the dorsal root ganglia

**DOI:** 10.1101/2024.01.20.576299

**Authors:** Lauren J. Donovan, Chelsie L. Brewer, Sabrina F. Bond, Aleishai Pena Lopez, Linus H. Hansen, Claire E. Jordan, Oscar C. González, Luis de Lecea, Julie A. Kauer, Vivianne L. Tawfik

## Abstract

Aging negatively impacts central nervous system function; however, the cellular impact of aging in the peripheral nervous system remains poorly understood. Aged individuals are more likely to experience increased pain and slower recovery after trauma. Such injury can damage vulnerable peripheral axons of dorsal root ganglion (DRG) neurons resulting in somatosensory dysfunction. One cellular mechanism common to both aging and injury is cellular senescence, a complex cell state that can contribute to the aged pro-inflammatory environment. We uncovered, for the first time, DRG neuron senescence in the context of aging and pain-inducing peripheral nerve injury in young and aged mice. Aged DRG neurons displayed multiple markers of senescence (SA-β-gal, p21, p16, IL6) when compared to young DRG neurons. Peripheral nerve injury triggered a further accumulation of senescent DRG neurons over time post-injury in young and aged DRG. These senescent neurons were dynamic and heterogeneous in their expression of senescence markers, p16, p21, and senescence-associated secretory phenotype (SASP) expression of IL6, which was influenced by age. An electrophysiological characterization of senescence marker-expressing neurons revealed high-firing and nociceptor-like phenotypes within these populations. In addition, we observed improvement in nociceptive behaviors in young and aged nerve-injured mice after treatment with a senolytic agent that eliminates senescent cells. Finally, we confirmed in human post-mortem DRG samples that neuronal senescence is present and increases with age. Overall, we describe a susceptibility of the peripheral nervous system to neuronal senescence with age or injury that may be a targetable mechanism to treat sensory dysfunction, such as chronic pain, particularly in aged populations.

## INTRODUCTION

Aging negatively impacts our physiology, with cellular-level changes influencing whole organ function. In the central nervous system (CNS), aging leads to increased risk of diseases, such as Alzheimer’s and Parkinson’s, which present with progressive neurodegeneration ultimately resulting in cognitive impairments.(1) How aging impacts the peripheral nervous system (PNS), and susceptibility to sensory dysfunction such as pain(2, 3), remains more elusive. Limited evidence suggests alterations in primary sensory neurons with age, with few studies that investigated the cellular mechanisms that may contribute to these changes.(4–7)

One aberrant state increasingly prevalent in all cells with aging is cellular senescence.(8, 9) Cellular senescence is a complex cellular state in which damaged or defective cells irreversibly cease cell division, resist apoptosis and cell death, and express a pro-inflammatory senescence-associated secretory phenotype (SASP)(10). Cellular senescence can occur in various tissues in the body, including the brain and spinal cord, in response to disease or injury in young and aged animals.(9, 11–13) With age, the clearance of senescent cells decreases, resulting in their accumulation and enhanced secretion of pro-inflammatory SASP factors, ultimately contributing to the progression of age-associated disease.(14–19) Importantly, elimination of these long-lasting senescent cells improves disease pathology underscoring their deleterious contribution to tissue function.(20, 21) Cyclin dependent kinase inhibitors p21^CIP1/WAF1^ (p21) and p16^INK4A^ (p16) are classic markers of senescent cells that function to halt cell-cycle and drive early and late-stage senescence programs, respectively.(22–24) While senescence has been extensively studied in mitotic cells, there is now evidence that post-mitotic cells can acquire senescent signatures such as expression of p21, p16, and associated SASP.(25–28) Intriguingly, human pyramidal, cortical, and myenteric neurons can express these same markers of senescence.(29–32) This suggests that human neurons also senesce in the aged or diseased nervous system, potentially staving off neuronal loss after aberrant cell cycle entry.(33–35) In Alzheimer’s disease, senescent neurons and glia are implicated in disease pathology and genetic or pharmacologic elimination of these senescent cells in mice can improve molecular and functional outcomes.(36, 37)

Primary sensory neurons, whose cell bodies reside in the dorsal root ganglion (DRG), are susceptible to damage of their peripheral axons in a variety of contexts including limb trauma or surgery. Injury-induced hyperexcitability of these neurons, mediated in part by chronic inflammation within the DRG, can contribute to long-lasting pain.(38–41) In particular, cytokine signaling exacerbates nociceptive neuron hyperexcitability through modulation of the Trpv1 ion-channel receptor.(42–46) Interestingly, the very inflammatory molecules that mediate this hyperexcitability are all common SASP factors released by senescent cells which may act as a potential source of these key pain-inducing molecules after injury.(47)

Using a comprehensive set of senescence markers including p21 and p16, senescence associated-β-galactosidase activity (SA-β-gal), and expression of SASP factor IL6, we identified senescent primary sensory neurons within the mouse lumbar DRG induced with age and after peripheral nerve injury. We further investigated the impact of senescence on intrinsic neuron excitability using electrophysiology, and assessed pain behaviors in young and aged mice after clearance of senescent cells using a senolytic agent. Finally, we characterized senescent phenotypes in human sensory neurons with age, providing a basis for further investigation of senescent sensory neurons in the DRG as an analgesic target in the context of age and nerve injury-induced pain.

## RESULTS

### Senescent sensory neurons increase with age in the mouse DRG

We first determined whether cellular senescence occurs in the peripheral nervous system by examining sensory neurons in the lumbar dorsal root ganglia (DRG). We screened for a classic marker of senescent cells, SA-β-galactosidase activity(10), in the DRG of young (11-16 weeks) or aged (20-24 months) male and female mice. We found an increase in SA-β-galactosidase activity in aged compared to young DRG, indicating increased senescence of cells in the DRG with age (Figure 1A). Based on morphology and size, a majority of DRG cells with SA-β-galactosidase activity were primary sensory neurons (Figure 1A).

**Figure 1.**
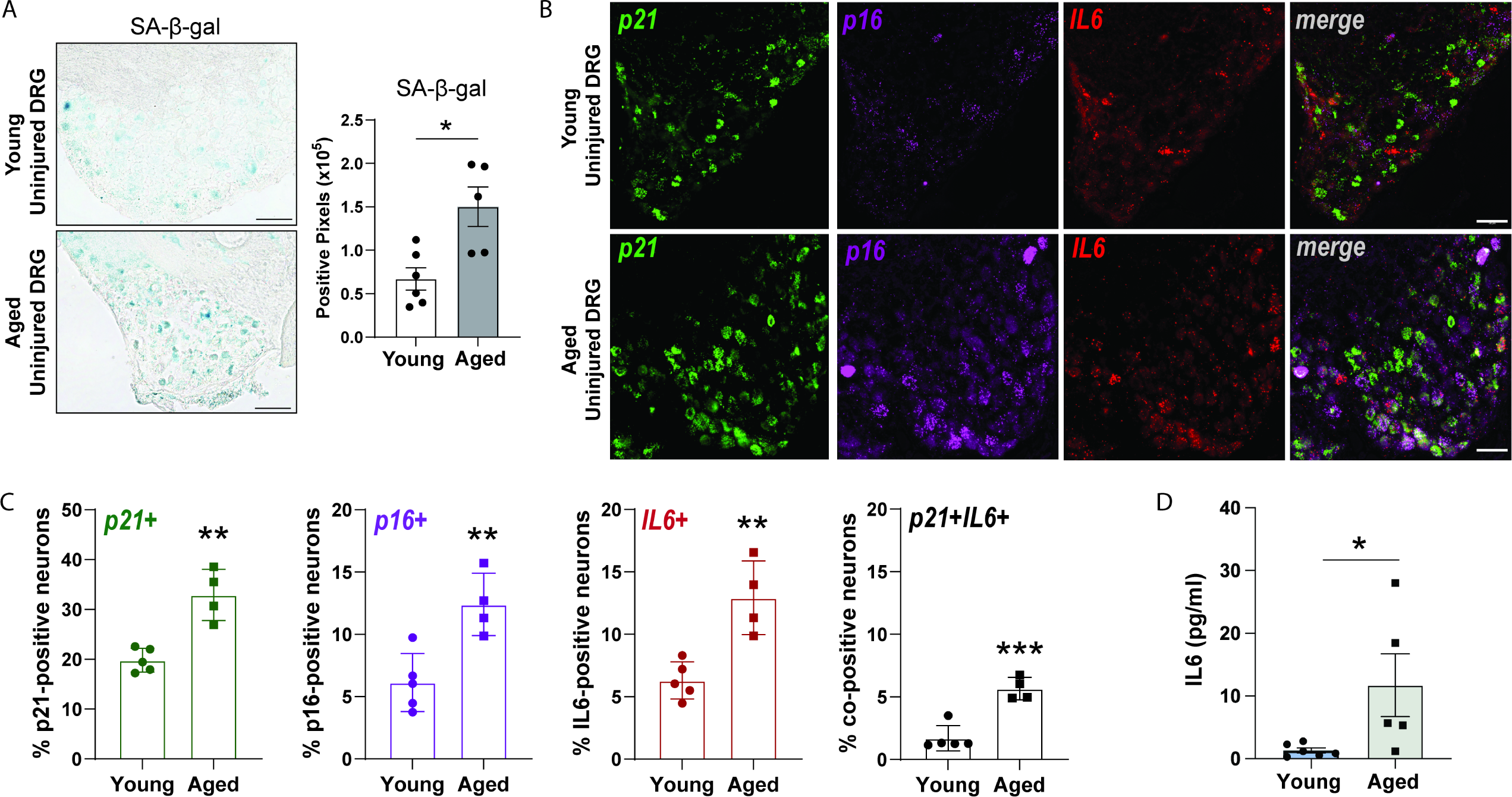
Senescent neurons accumulate with age in the mouse DRG. A. Representative images of SA-β-gal activity staining (*blue*) in the lumbar DRG of young and aged mice. Positive pixels detected were quantified (*right*) (n= 5-6 mice per group; two-tailed unpaired t-test, *p<0.05.). B. Representative RNAscope images for senescence markers p21 and p16 with SASP factor IL6 in whole DRG section. C. Quantification of neuronal expression of each marker or in combination expressed as a percent of total DRG neurons (n=4-5 mice per group, two-tailed unpaired t-test, *p<0.05, **p<0.01, ***p<0.001). D. Quantification of IL6 protein levels by ELISA assay in young or aged plasma (n=5-6 mice per group, two-tailed unpaired t-test, *p<0.05). Data are expressed as the mean ± SEM.

Although SA-β-gal activity is an indicator of senescence, its presence alone is insufficient for determining a senescent phenotype, which can be heterogeneous(10, 48). We therefore examined the RNA expression of two major drivers of senescence, *Cdkn1a* (p21^WAF1/CIP1^) and *Cdkn2a* (p16^INK4A^), in young and aged tissues by RNAscope. We detected a baseline expression of *p16* in 6% of DRG neurons in young mice (Figure 1B & C). Since the *Cdkn2a* transcript has two variants producing different protein products (p16^INK4A^ and p19^ARF^), we verified that the specific variant that produces p16^INK4A^ was in fact expressed by DRG neurons (Supplemental Figure 1). In aged mice, we detected significantly increased percentages of both *p21*+ and *p16*+ neurons compared to young mice, indicating enhanced senescence of DRG neurons with age (Figure 1B & C). As deleterious senescent cells are associated with pro-inflammatory SASP, we further co-localized senescence markers, *p21* and *p16*, with downstream SASP factor and cytokine *IL6*. The aged DRG displayed a significant increase in the number of *IL6*+ as well as co-positive *p21*+*IL6*+ neurons, when compared to the young DRG (Figure 1B & C). Additionally, enhanced IL6 protein levels were detected in the plasma of aged versus young mice (Figure 1D). These collective results indicate that senescent primary sensory neurons accumulate in the mouse DRG with age and express pro-inflammatory mediator and SASP factor IL6.

### Senescent neurons accumulate in the DRG following peripheral nerve injury

We next investigated whether direct injury to peripheral axons of primary sensory neurons would increase senescence of these cells in young adult mice. We performed spared nerve injury (SNI) in young mice (10-16 weeks old), in which two of the three distal branches of the sciatic nerve are transected and ligated with a suture(49) (Figure 2A). Lumbar L3/4 DRG were collected following injury to evaluate senescence (Figure 2A). Evaluating whole DRG by qPCR, we detected a significant increase in the RNA of senescence markers p21 and p16, and multiple SASP factors including IL6, IL1β, and Ccl2 in the ipsilateral SNI DRG at 3-weeks post-injury compared to controls (Figure 2B). To localize the cellular source of senescence marker expression in the DRG following SNI we next performed RNAscope for p21 and p16 transcripts in young male and female mice at several time points after SNI. The majority of *p21* and *p16*-expressing cells were neurons based on cellular morphology post-injury (Figure 2C). Neuronal expression of these senescence markers was quantified at acute (7-day), early chronic (3-week), and late chronic (7-week) time points following SNI to assess the induction and longevity of senescent neurons in the DRG post-injury (Figure 2A). Numbers of *p21*+ neurons increased significantly at the early 7-day time point in young injured mice and remained significantly increased throughout the time course when compared to the young uninjured mice (Figure 2D, *upper*). In contrast, we detected a gradual increase in the number of *p16*+ neurons over time following injury that became significant at 3-weeks post-SNI and peaked at the chronic 7-week post-injury time point compared to the young uninjured DRG (Figure 2D, *lower*).

**Figure 2.**
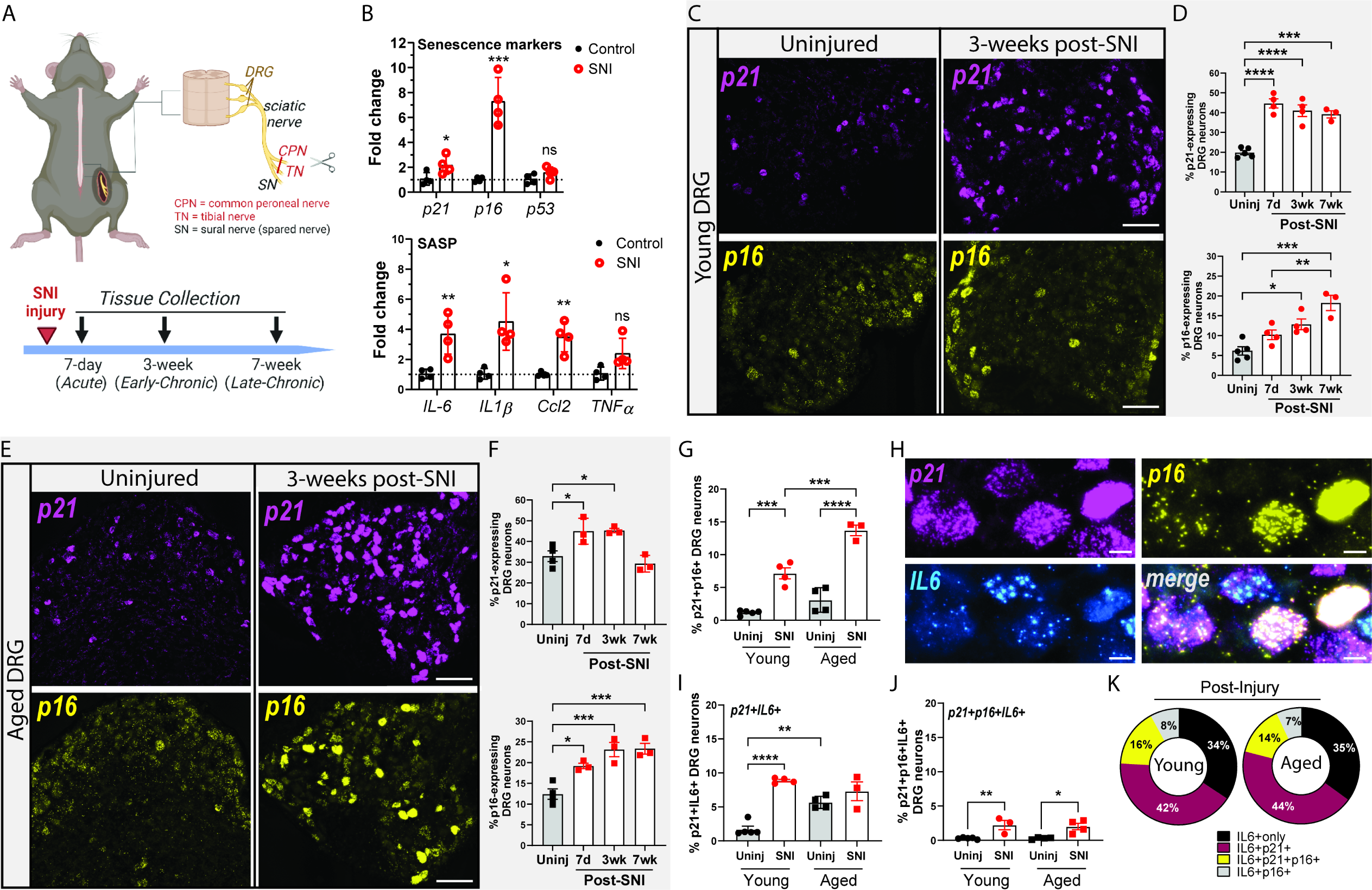
DRG neurons express senescence markers and SASP factors following peripheral nerve injury. A. Schematic of spared nerve injury (SNI) model in mice. Timeline over which DRG tissues were analyzed following SNI. B. qPCR of a panel of senescence markers and SASP factor gene expression from whole lumbar DRG in controls or following SNI (3-weeks post) in young mice (n=4 control; n=4 SNI young mice; two-tailed unpaired t-test, *p<0.05, **p<0.01, ***p<0.001). C. Representative RNAscope image of whole DRG section from uninjured or 3-weeks post-SNI in young mice. Scale bar 100µm. D. Quantification of the number of L3/4 DRG neurons expressing either *p21* (upper graph) or *p16* (lower graph) in young mice. (n= 3-5 mice per group/time-point; One-way ANOVA, *p<0.05, **p<0.01, ***p<0.001, ****p<0.0001). E. Representative RNAscope image of whole DRG section from uninjured or 3-weeks post-SNI in aged mice. Scale bar 100µm. F. Quantification of the number of L3/4 DRG neurons expressing either *p21* (upper graph) or *p16* (lower graph) in aged mice. (n= 3-4 mice per group/time-point; One-way ANOVA, *p<0.05, **p<0.01, ***p<0.001). G. Quantification of *p21*+*p16*+ co-expressing DRG neurons as a percent of total DRG neurons in uninjured and 3-weeks post-injury in young and aged mice. (n= 3-5 mice per group/time-point; One-way ANOVA, ***p<0.001, ****p<0.0001. H. Representative RNAscope images showing co-expressing senescence markers (*p21* and *p16*) with SASP factor (IL6). Scale bar 10µm. I. Quantification of *p21*+*IL6*+ co-expressing DRG neurons as a percent of total DRG neurons in uninjured and 3-weeks post-injury in young and aged mice. (n=3-5 mice per group; One-way ANOVA, **p<0.01, ****p<0.0001. J. Quantification of *p21*+*p16*+*IL6*+ triple positive DRG neurons as a percent of total DRG neurons in uninjured and 3-weeks post-injury in young and aged mice (n=3-5 mice per group; One-way ANOVA, *p<0.05, **p<0.01). K. Analysis of *IL6*-expressing DRG neuron population that co-express senescence markers *p21* and/or *p16* at 3-weeks post-injury in young and aged mice (n=3 mice per group). Data are expressed as the mean ± SEM.

Given that *p21* and *p16* expression can be induced in DRG neurons by nerve injury in young mice, we next tested whether senescence signatures were distinct in aged mice over the time course following injury (Figure 2A). Aged DRG neurons increased expression of *p21* and *p16* post-injury compared to aged uninjured controls (Figure 2E). Similar to young mice, we detected a significant increase in the number of *p21*+ neurons at 7-days and 3 weeks post-injury in aged mice, however numbers returned to aged uninjured levels by 7-weeks post-injury (Figure 2F, *upper*). In contrast to young mice, *p16*+ neurons were already significantly increased in the aged DRG by the early 7-day time point, and further increased and stabilized throughout the time course (Figure 2F, *lower*). In addition, there was a significant increase in neurons co-expressing both *p21* and *p16* in young and aged mice starting at 3-weeks after injury compared to their uninjured controls. Furthermore, the aged injured DRG displayed a significant increase in these *p21*+*p16*+ neurons compared to young injured DRG, suggesting the aged DRG have greater numbers of senescent cells transitioning to a late-stage p16-senescent state compared to young DRG (Figure 2G).

For a more robust detection of potential deleterious neuronal senescence within the DRG, we quantified the percentage of neurons that co-expressed any combination of *p21*, *p16*, and downstream SASP factor *IL6* (Figure 2H). Young mice accumulated significant numbers of neurons co-expressing *p21* and *IL6* in the DRG at 3-weeks post-injury (Figure 2I). While we detected increased baseline numbers of *p21*+*IL6*+ cells in aged uninjured mice compared to young, we did not find a further increase after injury in aged mice (Figure 2I), suggesting a heterogeneity of senescence marker induction after injury dependent on age. Triple positive *p21*+*p16*+*IL6*+ cells were increased in both young and aged mice after injury, albeit in low percentages out of total DRG neurons (Figure 2J). Finally, a majority (∼65%) of all IL6-expressing DRG neurons post-injury expressed either *p21*, *p16*, or both senescence markers in young or aged mice (Figure 2K). Collectively, these data support that nerve injury drives senescence in primary sensory neurons in the mouse DRG and that there is heterogeneity of senescent neuron phenotype. Further, these results suggest that senescent primary sensory neurons are a major and long-lasting cellular source of IL6 in the young and aged DRG post-injury.

### ATF3+ injured and neighboring non-injured neurons co-express p21 and p16 markers

We next hypothesized that neurons whose peripheral axons were injured by SNI would express senescence markers in the DRG. To test this, we co-labeled injured neurons, using ATF3 a marker of axonal injury(50, 51), with p21 and p16 at multiple time points post-SNI. First, we detected very few ATF3+ neurons in uninjured young or aged animals (Figure 3A). Post-injury, ATF3+ neurons increased to ∼44% in the young DRG and ∼35% in the aged DRG neurons, out of total DRG neurons (Figure 3A). Strikingly, the majority of all ATF3+ neurons co-expressed *p21* and/or *p16* in both young and aged mice at all time points post-injury (Figure 3B, *arrows* & 3C-D). We further detected an age-independent expansion of *p16*-expressing ATF3+ population at later time points following injury suggesting that injured neurons progressed into a late-stage p16-senescent state over time (Figure 3C-D). Further, not all *p21*- or *p16*-expressing neurons in the DRG were ATF3+ (Figure 3B *below asterisks*). The proportion of *p21* or *p16* expressing neurons that were ATF3-negative, either increased or remained stable over time after injury in young or aged mice (Figure 3E-F). These results demonstrate that non-injured neurons also senesce, potentially representative of “bystander” or “secondary” senescence in the DRG of nerve-injured mice.

**Figure 3.**
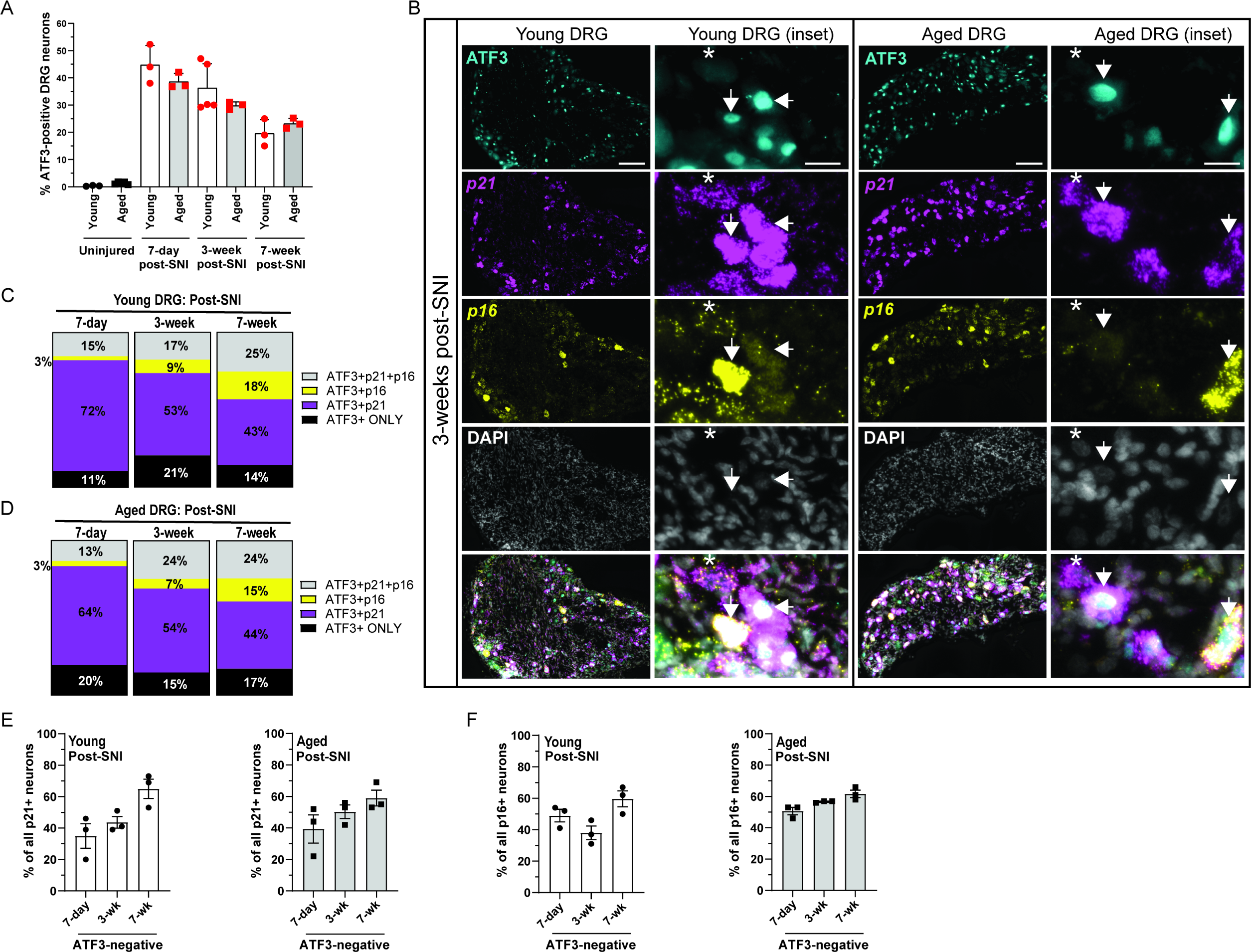
ATF3+ injured and neighboring non-injured DRG neurons express senescence markers after nerve injury. A. Quantification of number of ATF3+ neurons as a percent of total L3/4 DRG neurons in uninjured and multiple post-SNI time points in young and aged mice (n=3-5 mice per group/time-point). B. Representative images of dual immunohistochemistry/RNAscope labeling ATF3+ injured neurons (nuclear localized protein) and RNA puncta of p21 and p16 at 3-weeks post-SNI. Co-expression of ATF3 with *p21* and or *p16* (*arrows*). Asterisks represent ATF3-negative cells that express p21 and/or p16 senescence markers. C & D. Quantification of ATF3-positive neuron population that co-express *p21* and/or *p16* at multiple time points post-injury in young and aged DRG (*Young*: n=3-5 mice per time-point per group: 7-day: *n*=1,421 ATF3+ neurons, 3-week: *n*=1,056 ATF3+neurons; 7-week: *n*=523 ATF3+neurons; *Aged*: n=3 mice per time-point: 7-day: *n*=1,004 ATF3+ neurons; 3-week: *n*=983 ATF3+ neurons; 7-week: *n*=722 ATF3+ neurons). E. Quantification of ATF3-negative population co-expressing senescence marker *p21* at multiple time points post-injury in young and aged DRG (n=3 mice per group per time-point). E. Quantification of ATF3-negative population co-expressing senescence marker *p16* at multiple time points post-injury in young and aged DRG (n=3-5 mice per group per time-point). Data are expressed as the mean ± SEM.

### Trpv1+ nociceptors express senescence markers in the DRG

Individual subtypes of sensory neurons are tuned to respond to unique stimuli, are distinct in size, and vary in their expression of canonical markers(52). Therefore, to characterize the subtype(s) of primary sensory neurons that express senescence markers after nerve injury, we analyzed their cell diameters. Previously it was reported that the distribution of small-, medium-, and large-diameter DRG neurons do not change with age.(4) In our dataset, the majority of *p21*+*IL6*+ neurons measured in the range of 16-30µm, with a mean diameter of 22µm (+/- 5.49µm) in young and 20µm (+/- 4.96µm) in aged mice (Figure 4A). In comparison, *p16*+*IL6*+ neurons were slightly larger in diameter with a mean diameter of 29µm (+/- 5.51µm) in young and 24µm (+/- 5.96µm) in aged mice (Figure 4A). In either case, young or aged senescent neurons were rarely of large diameter (Figure 4A).

**Figure 4.**
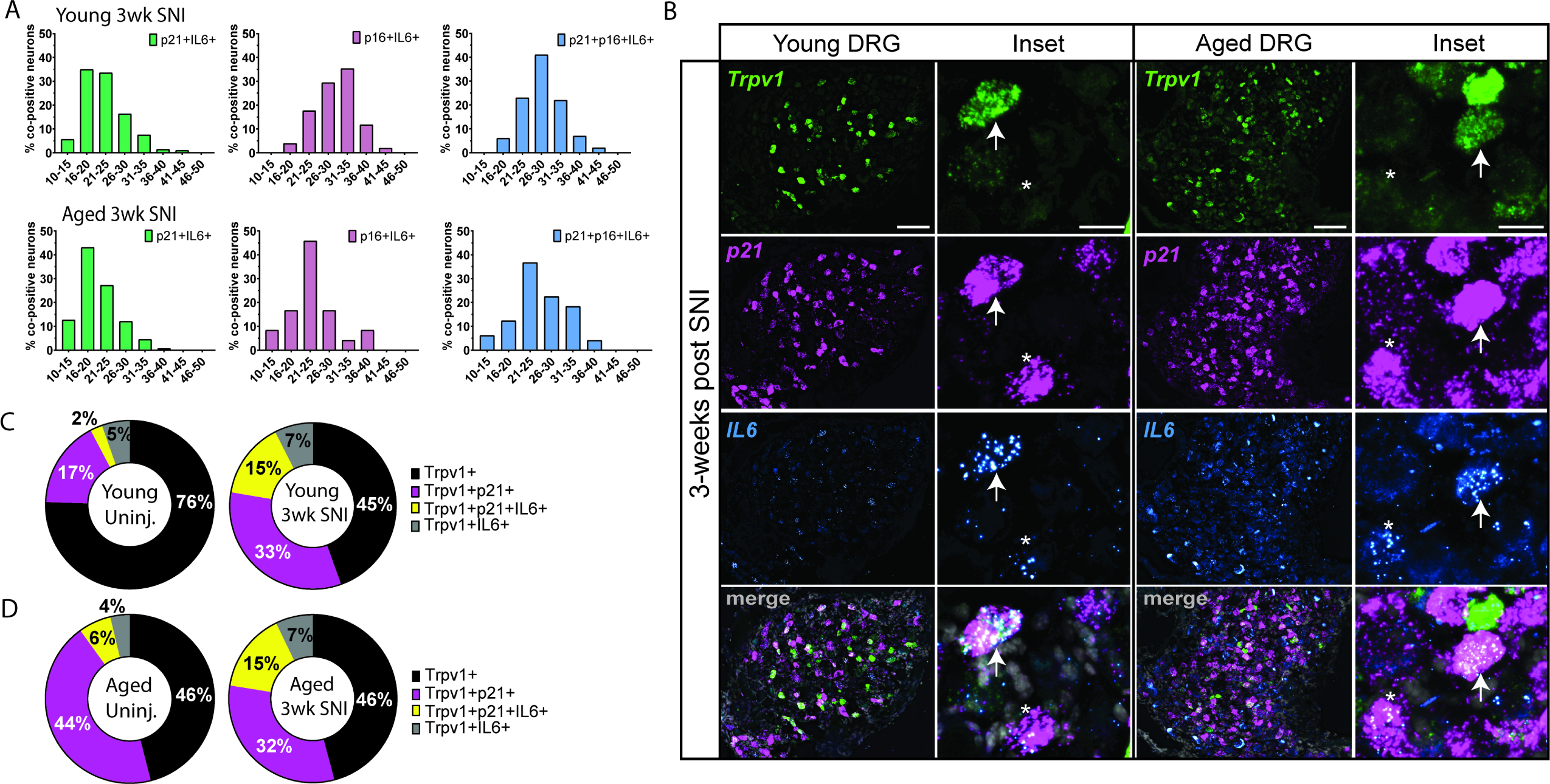
Trpv1+ nociceptors express senescence markers following nerve injury. A. Analysis of cell diameter (µm) of *p21*+*IL6*+, *p16*+*IL6*+, or *p21*+*p16*+*IL6*+ co-positive neurons in the DRG at 3-weeks post-nerve injury in young and aged mice (*Young*: *n*=215 *p21*+*IL6*+ neurons; *n*=51 *p16*+*IL6*+ neurons; *n*=102 *p21*+*p16*+*IL6*+ neurons; *Aged*: *n*=155 *p21*+*IL6*+ neurons; *n*=21 *p16*+*IL6*+ neurons; *n*=46 *p21*+*p16*+*IL6*+ neurons). B. Representative RNAscope images of young or aged DRG co-labeled for the ion-channel Trpv1, senescence marker p21, and SASP factor/cytokine IL6. Merged images also have DAPI overlay (grey). For *IL6*-signal, intense puncta signal with white center are positive neurons, while fainter/dull blue is background. Arrows: Trpv1+ senescent neurons, Asterisks: Trpv1-negative senescent neurons. Scale bars 100µm and 20µm (insets). C. Quantification of *Trpv1* neuron population and its co-expression with *p21* and/or *IL6* in young and (D) aged L3/4 DRG of uninjured (controls) and 3-weeks post-SNI (n=3 uninjured young mice, *n*=972 Trpv1+ neurons; n=3 SNI young mice, *n*=1,548 Trpv1+ neurons; n=4 uninjured aged mice, *n*=1,056 Trpv1+ neurons; n=3 SNI aged mice, *n*=1,292 Trpv1+ neurons).

Given that the majority of neurons expressing senescence markers were small diameter, we theorized that these senescent neurons may be Trpv1+, as this ion channel is widely expressed in small diameter neurons classified as nociceptive neurons.(39, 53) First, the majority of *Trpv1*+ neurons co-labeled with *p21* after injury, with fewer co-labeled with *p16* in young mice at 3-weeks post injury. To identify whether these *Trpv1*+*p21*+ neurons localized with downstream SASP factor IL6, we co-labeled these neurons at baseline (uninjured) and following SNI (Figure 4B). In young uninjured mice, we detected 19% *Trpv1*+*p21*+ neurons with only 2% co-expressing SASP factor *IL6*, which markedly increased after injury to 48% *Trpv1*+*p21*+ with 15% co-expressing *IL6* (Figure 4C). In aged uninjured mice, we detected 50% *Trpv1*+*p21*+ neurons with 6% co-expressing *IL6*, a 3-fold increase in *IL6*-expression compared to young uninjured DRG (Figure 4C & 4D). Similar to young mice after injury, we found an expansion of the aged *Trpv1*+ population co-expressing *p21*+ and *IL6*+ after injury compared to aged uninjured DRG (aged uninj: 6% vs aged SNI: 15%) (Figure 4D). Taken together these data support enhanced *Trpv1*+ nociceptor senescence with age and following injury, with an increased fraction that co-express *IL6* after injury in both young and aged mice.

### Senescent DRG neurons have unique physiological profiles

We next surveyed the electrophysiological properties of senescence marker-expressing neurons to characterize their excitability profile and gain additional insight into their functional contribution in the DRG. We used whole-cell patch-clamp in intact DRG preparations from young and aged mice with and without injury followed by single-cell PCR to detect the presence of *p16*, *p21*, and SASP factor *IL6*.

We recorded from 82 separate lumbar primary sensory neurons from 26 mice. Given the heterogeneity of DRG functional profiles, we employed dimensionality reduction using Uniform Manifold Approximation and Projection (UMAP, python implementation from https://github.com/lmcinnes/umap) based on 33 parameters collected during recordings, including firing properties, diameter, and intrinsic currents (Figure 5A). This analysis produced five discrete clusters as determined by a hierarchical density-based cluster algorithm (HDBSCAN; python implementation from https://github.com/scikit-learn-contrib/hdbscan; Figure 5B). We then analyzed the distribution of markers of senescence (p21-Figure 5C, p16-Figure D, and IL6-Figure 5E) within the clusters. Notably, the majority of *p16*-expressing neurons grouped in cluster 5 (83%, 5/6), a cluster that contained *all* of the neurons with high evoked firing phenotypes (defined as >100 action potentials [APs] fired during all current steps; Figure 5F, top heatmap), two of which were *p16*-positive (Figure 5F, top and lower left heatmap). Similarly, the majority of neurons expressing the IL6 were found in cluster 5 (Figure 5E) and included a high-firing phenotype neuron (13%, 1/8; Figure 5F, top and lower left heatmap). Cluster 5 also contained neurons with lower rheobase and higher hyperpolarization-activated current (Ih), both of which can contribute to increased excitability (Figure 5F, top and lower right heatmap). *p21*-expressing neurons, however, encompassed a larger population and were distributed throughout all clusters (Figure 5D & F). Given the basal expression of *p21* in a subset of young uninjured DRG neurons (∼20%) (Figure 1C), this distribution of *p21*-positive neurons across clusters indicates a mixed population of non-senescent and senescent neurons. Overall, the majority of high-firing neurons expressed at least one senescence marker (63%, 10/16; Figure 5F). Additionally, the senescence marker and/or SASP factor-expressing populations within our dataset contained neurons with wide APs (defined as AP half widths greater than 0.5 ms; *p16*: 67%, 4/6; *p21*: 72%, 26/36; *IL6*: 75%, 6/8; data not shown), suggesting they belong to nociceptor or C fiber low threshold mechanoreceptor classes, which express the Trpv1 channel, supporting our prior findings and hypothesis that Trpv1+ nociceptors senesce.(54) Together, these data suggest that senescence marker-expressing DRG contain populations of high-output DRG neurons and share physiological characteristics with nociceptors.

**Figure 5.**
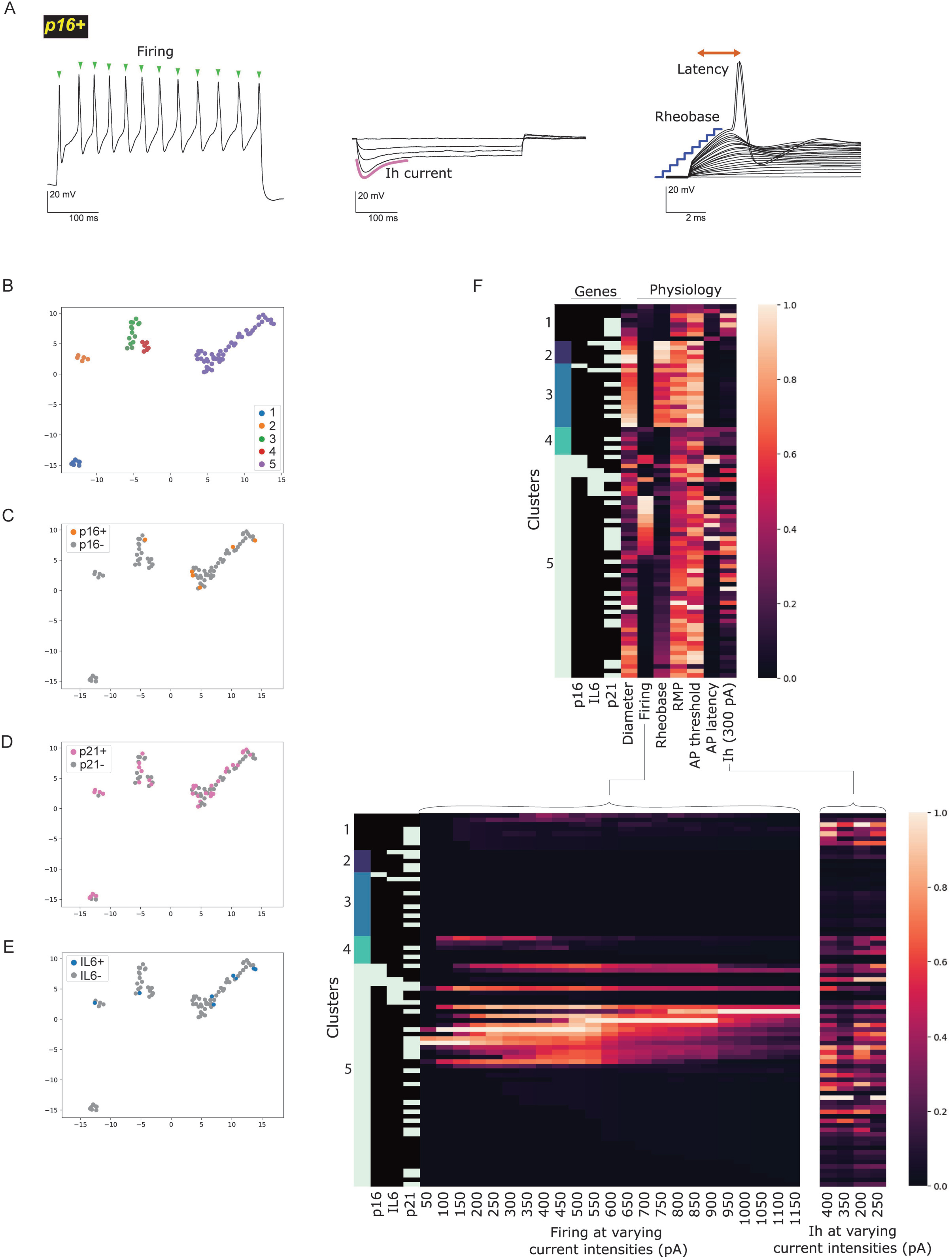
DRG neurons expressing senescence and SASP markers have unique functional profiles that include high-firing and nociceptor-like phenotypes. A. Representative traces from *p16*-expressing neurons demonstrating repetitive firing (*left*), hyperpolarization-activated current (Ih) presence (*middle*), and the firing parameters rheobase and AP latency (*right*). B. Clusters identified with the hierarchical density-based algorithm HDBSCAN after UMAP alignment of individual neurons constructed with diameter (range: 14– 41µm), firing properties and intrinsic currents. Discrete clusters are identified by color (1–5). (*n*= 82 recorded DRG neurons). C. The senescence marker p16 (Orange) and D. the SASP factor IL6 (*blue*) are mainly found in cluster five. E. The senescence marker p21 (*pink*) is widely distributed throughout the clusters. F. Heatmap depicting parameters from left to right as follows: Clusters (*cool gradient*), gene expression (*black*=no expression and *white*=expression), diameter and physiology parameters (*warm gradient*= higher normalized values in lighter reds and lower values in darker reds). Senescence marker *p16* and SASP factor *IL6* groups contain neurons with high-firing phenotypes (>100 APs fired during current steps), which is outlined over increasing depolarizing current steps (*lower left panel*). Ih current amplitude was also measured at decreasing hyperpolarizing steps (*lower right panel*).

### Pharmacological elimination of senescent cells ameliorates pain behaviors in aged mice

Given that nociceptive neurons within the DRG co-express senescence markers, *p21* and SASP factor *IL6*, after nerve injury in young and aged mice (Figure 4B, *inset asterisks*), and that senescent neurons contain high-output and nociceptor-like populations, we next hypothesized that clearance of senescent DRG neurons would improve pain behaviors induced by SNI. We tested senolytic ABT263 (Navitoclax), a peripherally restricted senolytic that promotes apoptosis of senescent cells.(55–57) We treated mice with ABT263 (100mg/kg, p.o. in vehicle) or vehicle (60% Phosol40PG, 30% PEG400, 10% Ethanol, p.o.) for 10-days starting at 3 weeks post-SNI, a time-point at which senescent neurons have accumulated in young and aged mice (Figure 6A). Treatment with ABT263 induced apoptosis of DRG neurons, as evidenced by a significant increase in the number of cleaved caspase-3 (CC3) positive (i.e. apoptotic) neurons in ABT263-treated mice compared to vehicle-treated controls (Figure 6B). In aged mice treated with ABT263, we observed a gradual improvement in mechanical allodynia up to at least ∼3-weeks following treatment, compared to vehicle-treated aged mice (Figure 6C). In addition, aged mice showed a significant improvement in weight bearing immediately (Day 16) and 3 weeks (Day 39) after senolytic treatment (Figure 6D). In young mice, there was a significant but transient reduction in mechanical threshold during the second 5-day treatment window of ABT263 when compared to vehicle treated mice (Figure 6E). In addition to this transient improvement in touch sensitivity, we observed sustained improved weight bearing on the injured hindlimb in young mice (Figure 6F). Sensory function of the contralateral (uninjured) hindlimb was not altered after application of senolytics in aged or young mice (Supplemental Figure 2A & B). Collectively, these data show senescent DRG neurons can be targeted by senolytics and that treatment can result in improved behaviors associated with pain outcomes in a nerve injury model and may be more effective in aged animals.

**Figure 6.**
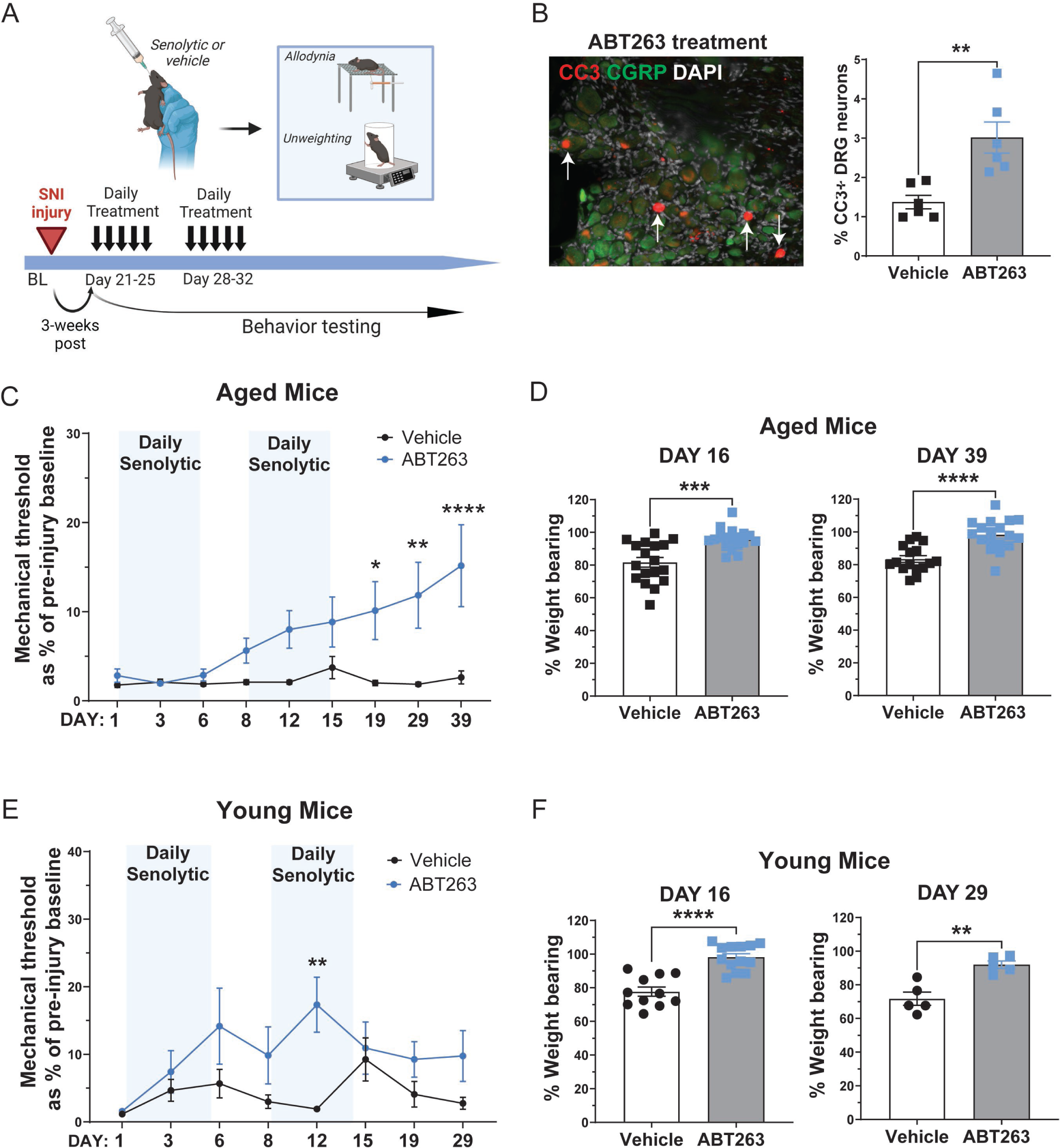
*in vivo* elimination of senescent neurons using senolytics alleviates pain behaviors after nerve injury. A. Schematic of treatment paradigm. Young and aged mice were treated with senolytic (ABT263, 100mg/kg) or vehicle for 10 days by oral gavage, starting at 3-weeks post-spared nerve injury (SNI). Mechanical allodynia and weight bearing were assessed during and after treatment. B. Senolytics induce apoptosis in the DRG. Representative image of CC3-immunohistochemistry to capture apoptotic cells after 5-day treatment with ABT263 (white arrows = CC3-positive neurons, other red fluorescent signal is background lipofuscin. Quantification of cleaved-caspase-3 (CC3) positive neurons in the DRG following treatment with vehicle or ABT263 for 5 consecutive days (n=3 male, n=3 female aged mice per treatment group, two-tailed unpaired t-test, **p<0.01). C. Aged mice were treated with ABT263 or vehicle (light blue indicates treatment window) and their mechanical allodynia thresholds were assessed (n=13 female, n=9 male vehicle-treated mice; n=14 female, n=13 male ABT263-treated mice, mixed-effects analysis, Šídák’s multiple comparisons test, *p<0.05, **p<0.01, ***p<0.001). D. Aged mice treated with ABT263 displayed improved weight bearing on injured limb compared to vehicle-treated mice at both Day 16 (n=10 female, n= 8 male vehicle-treated mice; n=9 female, n=9 male ABT263-treated mice, two-tailed unpaired t-test, ***p<0.001) and Day 39 post treatment start (n=9 female, n= 7 male vehicle-treated mice; n=9 female, n=9 male ABT263-treated mice, two-tailed unpaired t-test, ****p<0.0001). E. Young mice were treated with ABT263 or vehicle (light blue indicates treatment window) and their mechanical allodynia thresholds were assessed (n=12 male vehicle-treated mice, n=12 male ABT263-treated mice, mixed-effects analysis, Šídák’s multiple comparisons test, **p<0.01). F. Young mice treated with ABT263 displayed improved weight bearing on injured limb compared to vehicle-treated mice at both Day 16 (n=11 vehicle-treated, n=13 ABT263-treated male mice, two-tailed unpaired t-test, **p<0.01, ****p<0.0001) and Day 29 post treatment start (n=5 vehicle-treated, n=5 ABT263-treated male mice, two-tailed unpaired t-test **p<0.01). Data are expressed as the mean ± SEM.

### Human DRG neurons express senescence markers and accumulate with age

To validate the translational potential of senescent cell subpopulations as a target in humans, we assessed human DRG neuron expression of multiple senescence markers. As it is difficult to obtain human DRG tissues with confirmed injury/damaged nerves, we assessed whether senescence markers increase with age. We collected L4 DRG from a young (33yr) and an aged (65yr) female human donor and examined senescence marker expression by RNAscope. Similar to mice, human DRG neurons clearly expressed *p21* and *p16* (Figure 7A & B). When quantified, a greater percentage of aged human DRG neurons expressed either or both of these markers compared to young DRG neurons (*p21+*: 48% aged vs 24% young; *p16+*: 42% aged vs 36% young; *p21+p16+*: 40% aged vs 10% young) (Figure 7C). Further, greater numbers of IL6+ neurons were detected in aged compared to young human DRG (57% aged vs 36% young) (Figure 7D). Of this *IL6*-expressing neuronal population, aged DRG had an increased fraction that co-expressed either *p21* or *p16*, with a striking increase in triple positive cells, compared to young DRG (*IL6+p21+p16+:* 48% aged vs 23% young) (Figure 7E). Total neurons co-expressing these senescence markers were increased in aged compared to young DRG, demonstrating an increase in pro-inflammatory SASP-producing senescent cells with age (Figure 7F). Lipofuscin accumulation within cells can serve as an additional marker of lysosomal impairment and senescence.(58, 59) We observed that human DRG neurons had higher amounts of lipofuscin in cells, with many neurons >75% lipofuscin-filled, occluding the RNAscope signal and precluding further analysis of additional senescence marker expression (Supplemental Figure 3A). When quantified, we found an increased fraction of DRG neurons filled with lipofuscin with age in the human DRG, additionally indicative of senescence in the aged human DRG (9.5% aged vs 1.5% in the young) (Supplemental Figure 3B).

**Figure 7.**
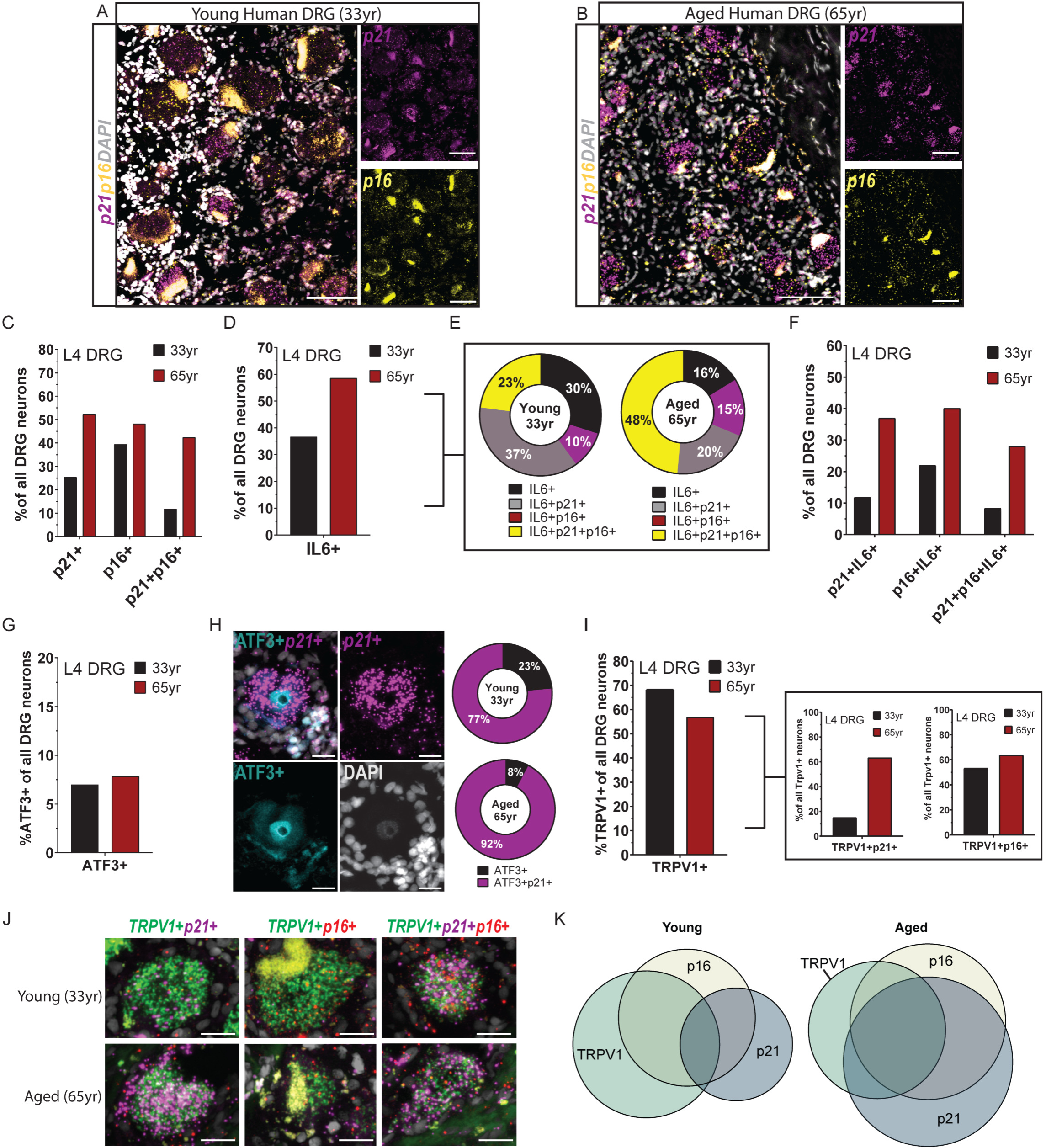
Human DRG neurons express senescence markers and SASP factor IL6 with age. A & B. Representative RNAscope images from young (33yr) or aged (65yr) human L4 DRG showing expression of *p21* and *p16* senescence markers (enlarged left images with DAPI, scale bar 100µm). The large globular signal present in both channels is auto fluorescent lipofuscin and not RNAscope signal (small puncta). C. Quantification of *p21*+, *p16*+, and co-positive *p21*+*p16*+ neurons in the young and aged human DRG as a percent of total DRG neurons. D. Quantification of *IL6*-expressing neurons as a percent of total DRG neurons. E. Analysis of *IL6*-positive neuron population and quantification of the co-expression of senescence markers *p21* and/or *p16*. F. Quantification of neurons co-expressing senescence markers *p21* and/or *p16* with *IL6* as a percent of total DRG neurons. G. Percent of DRG neurons that are ATF3-positive in young and aged human DRG. H. Example image depicting a single human neuron positive for ATF3 (nuclear-localized, immunohistochemistry) and *p21* (RNAscope). Scale bars 20µm. Analysis of ATF3-positive neuron population and quantification of the co-expression with p21 in young and aged human DRG (*right*, donuts) (*n*=64 young ATF3+ DRG neurons, *n*=54 aged ATF3+ DRG neurons). I. Total percentage of *TRPV1*+ neurons as a percent of total DRG neurons in young and aged human DRG. *Boxed right*, Quantification of the subsets of *TRPV1*+ neurons that co-express either *p21* or *p16* by RNAscope. J. Single human neurons showing co-expression of *TRPV1* with *p21* and/or *p16*. DAPI in grey. Scale bars 20µm. K. Venn diagram of human DRG neurons representing total numbers of DRG neurons that express *TRPV1*, *p16*, and *p21*. Aged DRG display a greater overlapping fraction of *TRPV1*+ neurons expressing either or both senescence markers *p21* and *p16 h*compared to young neurons.

We next asked whether injured (ATF3+) human sensory neurons existed in the young and aged human DRG.(60) We detected a limited number of ATF3+ neurons in either young or aged human L4 DRG tissues (Figure 7G). Strikingly, the majority of ATF3+ neurons co-expressed *p21*, with a greater percent expressing *p21* in aged (92%) compared to young (77%) human DRG (Figure 7H).

To determine which subtype(s) of sensory neurons were most susceptible to age-induced senescence in humans, we next analyzed neuron diameter. Similar to mouse DRG neurons, senescence marker-expressing human DRG neurons were largely of small diameter in both young and aged DRG (Supplemental Figure 4A & 4B). Finally, we verified whether human TRPV1+ nociceptors had increased expression of *p21* and *p16* in the aged versus young DRG. First, we detected ∼68% of all young DRG neurons expressed TRPV1 in L4 DRG and ∼55% were TRPV1+ in aged L4 DRG (Figure 7I, *left*). Of these TRPV1+ neurons, we found a higher fraction co-expressing either *p16* or *p21* in aged versus young DRG (Figure 7I, *right boxed* & 7J). Further, we observed a shift in TRPV1+ nociceptor co-expression with *p21* and *p16*, with a majority of TRPV1+ neurons (81%) expressing one or both markers in the aged DRG compared to young DRG (60%) (Figure 7J & K). These data collectively suggest that human DRG neurons senesce with age, including injured ATF3+ and TRPV1+ neurons, and that senescent neurons are a source of IL6 in the aging human DRG.

## DISCUSSION

Using several complementary approaches, we show for the first time that primary sensory neurons senesce with age and after peripheral nerve injury, and that targeting these neurons can improve sensory dysfunction. First, we discovered an age-associated increase in the neuronal expression of key senescence markers including *p21* and *p16*, as well as increased SA-β-gal activity, when comparing aged (20-24mo) to young (10-16wk) mouse DRG. This is in agreement with prior work showing that tissues throughout the body accumulate senescent cells with chronological age.(8) We validated this key concept in human DRG, providing the first evidence of senescence in human primary sensory neurons. Importantly, we further confirmed the senescence status of DRG neurons by detection of elevated levels of IL6, a well-established SASP component, co-expressed in *p21*-expressing neurons in aged compared to young mouse and human DRG. After SNI, which induces a direct injury to primary sensory neurons, both young and aged mice displayed a significant increase in neurons expressing senescence markers p21 and/or p16 when compared to uninjured controls. Senescence marker expression by DRG neurons was long-lasting, with an increased p16+ population over time post-injury in both young and aged DRG indicating irreversible, late-stage senescence.(22) In addition, unbiased clustering of senescence marker-positive neurons based on their electrophysiology parameters characterized these cells as high output DRG neurons. Lastly, DRG neurons targeted *in vivo* by a senolytic ABT263 (Navitoclax) resulted in improved mechanical allodynia and weight bearing on the injured hindlimb.(55, 56)

The concept of neuronal senescence is an emerging one, as the classic hallmark senescence feature of cell-cycle arrest is generally thought to be absent in post-mitotic cell types.(20) However, multiple other hallmark features of senescence can be present in post-mitotic cells including expression of cyclin dependent kinase inhibitors (p21, p16), SA-β-gal activity, DNA damage/oxidative stress markers, and SASP component expression.(61) Such findings in the CNS prompted a new term ‘postmitotic cellular senescence’ (PoMiCS) to complement this observed phenotype, with SASP being an important detrimental feature of post-mitotic senescent cells.(27, 61, 62) Why would post-mitotic cells such as neurons upregulate cyclin-dependent kinase inhibitors such as p21 or p16? In neurodegenerative conditions, neurons aberrantly re-enter the cell cycle in response to cellular stress, which can lead to cell death of damaged neurons.(34, 63–68) This may explain why CNS neurons senesce in the first place; expressing cyclin-dependent kinase inhibitors (p21 and p16) to halt the cell cycle, induce senescence, and ultimately stave off apoptosis to preserve neuronal numbers.(20, 33, 61) Depending on the context, however, it may not be beneficial to preserve neurons that are damaged; acute or transient senescence may be beneficial while chronic or persistent senescence may result in organ dysfunction.(69) We speculate that primary sensory neurons undergo a similar stress response mechanism of cell-cycle re-entry upon nerve injury resulting in post-mitotic cellular senescence. Indeed, DRG neurons can even re-enter the cell cycle in response to other stressors such as growth factor restriction and chemotherapy.(70, 71)

The spared nerve injury (SNI) model of neuropathic pain used in this study provides a unique opportunity to study neuronal senescence with spatiotemporal control. The abrupt nature of axonal injury results in a time-locked induction of DRG neuronal senescence. This is in contrast to investigating senescence in slowly progressing disease models such as Alzheimer’s.(36, 37) Additionally, the direct injury to the primary sensory neuron in our model allows for the study of discrete populations of senescent cells, in contrast to models of traumatic brain and spinal cord injury(72–74) that damage multiple cell types at once. Using this spatially restricted and injury-triggered induction of senescence we were able to track neuronal senescent phenotypes over time post-nerve injury and identify characteristics distinct during aging in both injured and uninjured neurons. Over the post-injury time course, we captured a phenotypic shift (i.e. p21-to-p16 expression) of DRG neurons. Specifically, in the young DRG, we detected an early spike in p21+ neurons, that declined over time, while the p16+ neuron population gradually increased over time. Similarly in human fibroblasts, p21 expression was found to decrease after senescence was achieved, suggesting subsequent upregulation of p16 expression by these cells as essential to maintain senescent-cell-cycle arrest.(22) Interestingly, this p16+ population increase was more robust at earlier time points post-injury in the aged DRG, indicating that aged neurons progress to late-stage senescence quicker compared to young neurons. Both injured and non-injured neurons expressed senescence markers over time post injury, suggesting this model captures a heterogenous population of 1) primary senescence, induced directly by the cell stressor (axon injury), and 2) secondary “bystander” senescence, induced by paracrine action of SASP-producing primary senescent cells.(23, 75–77) In support of this, the population of senescent neurons that were ATF3-negative (non-injured) increased over time in young and aged animals post-injury. While we did not find a difference in percentage of ATF3-positive neurons with age alone, we did find an overall increase in senescent neurons with age, suggesting that age-associated senescence occurs independent of injury. These same findings were further validated in human DRG tissues. These observations together support the identification of at least three populations of senescent neurons in this mouse injury model: age-induced, injury-induced, and secondary neuronal senescence; making it a unique model to study neuronal senescence heterogeneity. Further investigation into the deleterious impact on DRG tissue function by any of these senescent sub-populations will guide strategies to target appropriate cells for therapeutic purposes.

In this study, we found most IL6-expressing DRG neurons were senescence marker-expressing in young or aged animals after injury, indicating that senescent DRG neurons are a major source of IL6 in the DRG at three weeks post-nerve injury. Release of IL6 in the DRG can impact excitability of nociceptors via ion channel modulation, resulting in pain.(78–83) Production of cytokines such as IL6 in the DRG post-injury has been largely attributed to infiltrating immune and glial cells, namely macrophages and Schwann cells, which subsequently contribute to neuronal sensitization.(84–87) Further, while it has been reported that IL6-expressing neurons exist in the DRG following nerve injury (88), their function and the mechanism by which IL6 expression is induced remains unknown. We postulate senescence as a novel mechanism by which neurons produce IL6 as part of their SASP profile, which in turn acts on DRG neurons in an autocrine and/or paracrine manner,(78) ultimately resulting in hyperexcitability and pain. Intriguingly, the expansion of Trpv1+ nociceptors co-expressing both p21 and IL6 with age and after injury in mice and with age in human DRG provided evidence that this particular population, highly implicated in pain sensation, can be impacted by such cytokine production. Clinical studies support aberrant DRG excitability contributing to pain, evidenced by the effectiveness of peripheral anesthetics(89) or nerve blockade(90) in patients with chronic pain. Our finding that senescent neurons contain populations with high-firing or nociceptor-like characteristics supports the hypothesis that these neurons may directly contribute to hypersensitivity.

To evaluate the functional contribution of cellular senescence to sensory dysfunction we employed the senolytic ABT263, a small molecule drug that selectively targets senescent cells by acting on Bcl-2 anti-apoptotic pathways.(55, 56) Three other studies employed such an approach demonstrating that administration of senolytics in a model of chemotherapy-induced peripheral neuropathy(91) or nerve injury(12, 92) can improve behavioral outcomes, however DRG neurons were not implicated. Preserving sensation by eliminating sensory neurons may sound contradictory, however, correct timing of elimination (i.e. before too many cells enter senescence) may enable preservation of overall primary sensory neuron numbers. For example, elimination of senescent cells enabled overall survival of the many unaffected retinal ganglion cells in experimental ocular hypertension as well as cortical neurons in the context of AD pathology.(37, 93) For senescent neurons to be a specific and viable target, full characterization of the deleterious senescent cell population is needed. To this end, we are currently working on additional studies to better characterize senescent DRG neurons using single-cell sequencing to fully capture the phenotype and deleterious SASP profile of this heterogeneous population.

Given the extensive literature implicating cellular senescence in a variety of age-related pathology, our data extend the importance of this process to the peripheral nervous system. We further identify a novel cellular source of the pain-producing cytokine IL6 in the DRG, senescent primary sensory neurons, which are a long-lasting cell population after nerve injury. Overall, we describe a susceptibility of the peripheral nervous system to neuronal senescence with age or injury that may be a targetable mechanism to treat sensory dysfunction, such as chronic pain, particularly in aged populations.

## METHODS

### Animals

All mice were housed 2-5 per cage maintained on a 12-hour light/dark cycle in a temperature-controlled environment with *ad libitum* access to food and water. Young male and female mice used in this study were 10-16 weeks old, wild type C57BL/6J mice (Jax stock #00664). Aged male and female mice used in this study were 19-24months old, wild type C57BL/6JN mice (NIA aged rodent colony). Aged mice were pre-screened for abnormal masses and cataracts and otherwise were included in the study only if they appeared healthy.

### Human samples

Human lumbar L4 DRG tissues were obtained from one white female donor (age 33) who died from head trauma and one white female donor (age 65), who died from stroke.

### Study approval

All procedures were approved by the Stanford University Administrative Panel on Laboratory Animal Care and Institutional Animal Care and Use Committee in accordance with American Veterinary Medical Association guidelines and the International Association for the Study of Pain. Human post-mortem DRG was obtained in collaboration with Donor Network West and received Stanford University Institutional Review Board for human subjects exemption.

## METHOD DETAILS

### Spared Nerve Injury

To perform SNI surgery(49), mice were anesthetized with isoflurane and a small incision is made over the left thigh and blunt dissection is performed through the biceps femoris muscle in order to expose the sciatic nerve and its three branches (common peroneal, tibial, and sural nerves). The common peroneal and tibial nerves are then ligated using an 5-0 nylon suture (ETHILON^TM^ ref#1668G) and these nerves are then axotomized using small-sized spring scissors. The sural nerve is left intact (the “spared nerve”). The incision is then closed with surgical staples. Following surgery, mice are monitored for the study period, which varies from 1 day to 16 weeks depending on the time point of interest. Controls used for qPCR experiments, were sham surgery in which an incision was made followed by opening of muscle to reveal the nerve, without touching the nerve, followed by closure.

### Senolytic Administration

ABT263 (Navitoclax) (Med Chem Express, Cat. No.: HY-10087) was dissolved in 60% Phosol40PG, 30% PEG400, 10% Ethanol at a concentration of 12.5 mg/mL using brief water bath sonication and vortexing. Young and aged mice were briefly anesthetized with isoflurane before they were dosed by oral gavage (p.o.) at 100 mg/kg daily for 5 days, with a 2-day rest period, followed by a second 5-day daily dosing.

### Quantitative Real Time RT-PCR (qPCR)

Whole DRG tissues were collected, homogenized using a 1mL glass homogenizer, (PYREX® No. 7724-1) and placed in TRIzol Reagent (Invitrogen, Cat#15596018). RNA was isolated using miRNeasy® Mini kit (Qiagen, Cat #217004). The concentration and purity of RNA samples were determined using NanoDrop 2000 (Thermo Fisher Scientific). RNA was reverse transcribed using SuperScript^TM^ VILO^TM^ cDNA Synthesis Kit (Cat#11754-050). qPCR analysis was performed with PowerUp SYBR Green Master Mix (Thermo Fisher Scientific, Cat#A25741) and run on an Applied Biosystems 7900HT or on an Applied Biosystems StepOnePlus. Appropriate no reverse-transcriptase and no template controls were used for each 384-well PCR reaction. The cycle conditions were as follows: 50°C for 2 min, 95°C for 2 min, then 40 cycles of 15 s at 95°C, 1min at 60°C. Dissociation analysis was performed at the end of each run to ensure specificity. Relative quantification of gene expression was performed via 2^-ΔΔC(T)^ method.(94)

### Tissue preparation

Mice were anesthetized using Pentobarbital (Vortech Pharmaceuticals, NDC 0298-9373-68, 150mg/kg in 0.9% saline) and transcardially perfused with 5mL 1XPBS followed by 30mL 10% formalin solution (ThermoFisher). Lumbar DRG tissues were dissected and placed temporarily in RNAlater solution at RT (ThermoFisher), then frozen in O.C.T. Compound (Sakura Finetek, Inc., Cat#4583) on dry ice, and stored at -80°C. Mouse DRG was sectioned at 14µm and mounted onto SuperFrost Plus glass slides, dried for 1hr, and stored at −80°C until histology protocols performed. Human lumbar DRGs were obtained from organ donors and no identifying information was shared with the researchers. Nerve endings were trimmed and tissues were flash frozen immediately on dry ice and stored in screw cap 15mL conical tubes and stored at −80°C. DRGs were slowly embedded in OCT to avoid thawing and sectioned at 20µm onto SuperFrost Plus glass slides and stored at −80°C until histology protocols performed.

### Fluorescent in situ hybridization (RNAscope® Multiplex)

Fluorescent *in situ* hybridization using the RNAscope® Multiplex V2 Kit (ACD, Cat#323100) was performed in combination with immunohistochemistry to detect the RNA of senescence markers, cytokine, and DRG subtype markers (p21 (*Cdkn1a*), p16 *(Cdkn2a*), *IL6*, *Tprv1*) and protein markers (ATF3), respectively. Briefly, DRG tissues were isolated and processed as described above. Dorsal root ganglion (DRG) sections (14µm) were mounted on glass slides and dried 1-hr at room temperature and transferred to −80°C for storage. On Day 1 of RNAscope, slides were submerged in 10% formalin and incubated for 20 min at 4°C. Slides were washed with 1XPBS and dehydrated in EtOH as described in the ACD RNAscope user manual (UM 323100). Sections were incubated for 10min in RNAscope® Hydrogen Peroxide solution, washed in Millipore water, and incubated in RNAscope® ProteaseIV for 2min (human) or 5min (mouse) at room temperature. Slides were incubated with appropriate RNAscope probes (Mouse probes: Mm-IL6-C1, Cat#315891, Mm-Cdkn1a-C2, Cat#408551-C2, Mm-Cdkn2a-C3, Cat#411011-C3, Mm-Cdkn2a-tv2-C2, Cat#447491, Mm-Trpv1, Cat#313331-C1 and -C3; Human probes: Hs-TRPV1-C1, Cat#415381, Hs-CDKN2A-C2, Cat#310181-C2, Hs-CDKN1A-C3, Cat#311401) at 40°C in a Hybez^TM^ II oven (ACD) for 2hrs and stored overnight in 5X SSC buffer at room temperature. On Day 2, according to the ACD RNAscope user manual, slides were incubated in Amp1, Amp2, and/or Amp3 solutions followed by HRP-C1, HRP-C2, and/or HRP-C3 as appropriate. In each round, Opal^TM^ (1:1000, Akoya Biosciences Inc., OPAL 520 Cat#FP148700KT, OPAL 570 Cat#FP1488001KT, OPAL 690 Cat#FP1495001KT) or TSA Vivid^TM^ (1:1000, ACD Bio., TSA Vivid 520 Cat#323271, TSA Vivid 570 Cat#323272, TSA Vivid 650 Cat#323273) dye reagents and HRP Blocker. Negative control probes (ACD, #321838) were used to assess background levels of RNAscope signal.

### Immunohistochemistry

For immunohistochemistry performed immediately following RNAscope protocol (dual labeling), slides were first washed in 1XPBS and blocked (10% Normal Donkey Serum, 0.3 % Triton-X 100, in PBS) for 1hr at room temperature. Slides were incubated with Rabbit anti-ATF3 (1:200, Novus Bio, Cat#NBP1-85816) in 1% blocking solution in 1XPBS at 4°C overnight. Slides were washed 3X in 1XPBS for 5 min each, incubated with AlexaFluor secondary antibodies (1:1000, Donkey anti-rabbit-A488, LifeTechnologies, Cat# A21206), and mounted with Fluoromount G with DAPI (ThermoFisher, Cat#00-4959-52). For cleaved caspase-3 staining, mice were transcardially perfused as described and DRG tissues were extracted and frozen in OCT. Slides were then blocked (5%Normal Donkey Serum, 0.3 % Triton-X 100, in PBS) for 1 h at room temperature. Rabbit Cleaved-caspase-3 primary antibody (1:200, Cell Signaling Technology, Cat#9661) was incubated overnight at 4°C. Slides were incubated with secondary antibody, (1:1000, Donkey anti-rabbit Alexa-555, LifeTechnologies, Cat#A31572) for 2hr in the dark, washed, and mounted with Fluoromount G with DAPI.

### SA-β-galactosidase activity assay

Mice were perfused with PBS. L3-L5 dorsal root ganglion (DRG) was extracted, rinsed in RNAlater and PBS, and then mounted onto OCT. DRG were sectioned at 14mm onto glass slides. Slides were removed from the freezer, and 1X of fixative solution provided by Senescence β-Galactosidase Staining Kit (Cell Signaling Technology kit, Cat#9860S) was added to the slides for 15 minutes. Slides were rinsed in PBS, and a wax barrier was drawn around the edges of the sections. Fresh β-Galactosidase Staining Solution staining solution at pH 6.1 was added to the slides and then incubated at 37 °C for 22 hours. β-Galactosidase Staining Solution was removed and slides were rinsed twice in PBS and twice in distilled water before mounting and imaging. Imaging was done in FIJI ImageJ, where the area of the DRG neurons was outlined, and the percentage of positive β-Gal signal in the area was acquired. Sections (5–10) were analyzed per mouse. The total area analyzed per group was not significantly different from each other (data not shown).

### Neuron diameter analysis

Fluorescent TIFF images taken from RNAscope experiments that labeled *p21*, *p16*, and *IL6* RNA were used to measure diameters of neurons in both mouse and human DRG. Cells were individually labeled and categorized for the co-expression of markers *p21*, *p16*, with *IL6*. Using Fiji software, the scale (µm) was appropriately set based on objective used in image. The longest end-to-end cell diameters, with line placed through the center of each neuron, were drawn using the line segment tool. The line segment was then measured using ‘Measure’ as an output in µm unit and recorded. Neurons per category of RNA co-expression were then binned into 10µm segments and the percentage of neurons that fell into each µm bin were displayed as a percentage of total neurons analyzed in each subgroup.

### ELISA assay

1mL syringes were coated with heparin and used to withdraw ∼0.8ml of blood from mice anesthetized mice. To extract the plasma, the blood was centrifuged at 2000 RCF at 4°C for 10 minutes. The supernatant was removed, aliquoted, and stored at -80°C until used. IL-6 plasma concentration levels were assessed by following the manufacturer’s instructions (ThermoFisher, Cat#KMC0061). Samples were tested in duplicate and diluted 1:2. The final concentration was corrected for the dilution factor. A VERSAmax tunable microplate reader (Molecular Devices) was used to calculate optical density (OD) values at 450 nm. The data was analyzed with Boosterbio’s 5PL regression model and subtracting the blank well’s OD value from the sample’s OD values (https://www.bosterbio.com/biology-research-tools/elisa-data-analysis-online).

### Electrophysiology

Animals were deeply anesthetized with a ketamine/xylazine bolus (0.2 ml of 37.5 and 0.25 mg/ml in sterile saline, respectively). Mice were transcardially perfused with a sucrose-based dissection solution (containing in mM: 250 sucrose, 2.5 KCl, 25 NaHCO3, 1 NaH2PO4, 6 MgCl2, 0.5 CaCl2, and 25 glucose). All extracellular solutions in contact with live tissue were bubbled with a 95% O2/5% CO2 gas and chilled on ice, then decapitated. The vertebral column and sciatic nerves were isolated and placed in dissection solution. The DRG with attached nerves and dorsal roots were manually freed from the bone and muscle and stripped of epineurium. The tissue was transferred to collagenase (1 mg/ml in dissection solution) to incubate for 30 minutes at 35°C to allow for digestion of the perineurium. DRG recordings were performed in a chamber (RC-26GLP; Warner Instruments) within an upright microscope with platform (Nikon Eclipse FN1) and secured with a platinum wire-based anchor. The nerve end of the preparation was secured in a suction electrode attached to a stimulus isolator (A365, World Precision Instruments). Neurons were visualized with infrared differential interference contrast illumination. Recordings were done with patch pipettes pulled (P-97; Sutter Instruments) from single-filament borosilicate glass capillaries (1.5 mm OD, 1.1 mm ID; Sutter Instruments) with resistances from 5–8 MΩ, and internal patch solution as follows (in mM): 120 potassium gluconate, 20 KCl, 165 2 MgCl2, 2 Na2ATP, 0.5 NaGTP, 20 HEPES, 0.5 EGTA, pH adjusted to 7.2–7.3 with KOH. Signals were amplified (Multiclamp 700B; Molecular Devices), digitized (Digidata 1440A; Molecular Devices), filtered with a 4 kHz Bessel and sampled at 10 kHz (pClamp 10.6 software; Molecular Devices). Liquid junction potentials (-14 mV) were corrected for (JPCalc software, P. Barry, University of New South Wales, Sydney, Australia; modified for Molecular Devices). In current clamp, gap-free recordings were taken for 2 minutes (to measure spontaneous firing and membrane potential), then depolarizing current steps were applied from resting membrane potential in 50 pA steps (to measure rheobase, action potential [AP] threshold and AP latency) and finally, stepwise current pulses were injected from resting membrane potential to measure evoked firing frequency and Ih current (-300–1400 pA in 50 pA steps, 800 ms duration). Ih is reported as current density, which was computed as Ih current (amplitude in pA)/cell size (diameter in µm). Following recordings, images were taken of the neuron to estimate size (the average of two separate diameter measurements), and the cytoplasm was aspirated into the patch pipette for subsequent polymerase chain reaction (PCR). PCR was performed using primers for p16 (Mm.PT.58.42804808; IDT), p21 (Mm.PT.58.5884610; IDT), IL6 (Mm.PT.58.10005566; IDT), GFAP (To determine glia presence in sample; Mm01253033_m1; Thermo Fisher) and Tubb3 (To confirm neuronal tissue was sampled; Mm.PT.58.32393592; IDT) in combination with TaqMan™ Gene Expression Master Mix (Cat. #4369016; Thermo Fisher). Samples were then subjected to real-time PCR with the same primers and the SuperScript™ III One-Step RT-PCR System with Platinum™ Taq DNA Polymerase (Cat. #12574018; Thermo Fisher) to determine gene presence or absence in the sample. Of note, glial fibrillary acidic protein (GFAP) was present in several of our samples, thereby precluding us from determining the definitive source of p16, p21 or IL6 (i.e. DRG neurons or glial cells); nonetheless, our results reflect the state of the microenvironment in which the DRG was sampled. We intentionally recorded from mainly small-to medium-sized DRG to increase the chances of sampling senescence marker-expressing neurons. Uniform Manifold Approximation and Projection (UMAP; python implementation from https://github.com/lmcinnes/umap) was performed to use machine learning to integrate and connect the high-dimensional neuronal parameters (33) in low-dimensional 2D space. Training was performed using the train_test_split function from Scikit-Learn over 1000 epochs. UMAP hyperparameters were as follows: number of neighbors=5, minimum distance=0.82, local connectivity=2, random state=42. Clusters were then estimated via the hierarchical density-based clustering algorithm HDBSCAN (python implementation from https://github.com/scikit-learn-contrib/hdbscan/blob/master/docs/index.rst) with the following parameters: minimum cluster size=4, cluster selection epsilon=5, cluster selection method=’eom’ or Excess of Mass. Parameters were normalized for heatmap visualization using the following equation: (p - min(p)) / (max(p) - min(p)) where p is a vector containing all measurements of a given parameter.

### Mechanical nociception assays

To evaluate mechanical reflexive hypersensitivity, we used a logarithmically increasing set of 8 von Frey filaments (Stoelting), ranging in gram force from 0.007 to 6.0 g. These were applied perpendicular to the plantar hindpaw with sufficient force to cause a slight bending of the filament. A positive response was characterized as a rapid withdrawal of the paw away from the stimulus filament within 4 s. Using the up-down statistical method,(95) the 50% withdrawal mechanical threshold scores were calculated for each mouse and then averaged across the experimental groups. Mechanical nociception testing was performed at 3-weeks post SNI at Days: 1, 3, 8, 12, 15, 19, 29, 39 post-senolytic treatment start.

### Unweighting

An incapacitance device (IITC Life Science) was used to measure hindpaw unweighting. Mice were placed in the plexiglass apparatus with a ramp with the hindpaws resting on separate metal scale plates. Measurements were taken when the hindpaws were supporting the weight of the mouse with forepaws on the ramp. The duration of each measurement was 4-6 s, and 6 consecutive measurements were taken at 60 s intervals. Six readings were averaged to calculate the bilateral hindpaw weight-bearing values. Unweighting was measured post-senolytic or vehicle treatment in SNI young and aged mice. The calculation of weight bearing on the injured hindlimb was as follows: 2*(L)/(L + R)*100 to get percent weight bearing on injured (L: *left*) hindlimb.

### Imaging/Image analysis

All imaging was performed using a Keyence BZ-X810 fluorescent microscope (Keyence) with a 40X objective (mouse DRG) or a 20X objective (human DRG). Eight-12 DRG sections were imaged per mouse and 3-5 DRG sections per human sample. Images were saved as stitched and full focus TIFF files and analyzed. All images were similarly adjusted for brightness and contrast per experiment, with no additional alterations made to the image. Lipofuscin signal was defined by strong autofluorescence signal across all channels (488, 550, 647).

### Quantification and statistical analysis

Measurements of Cohort sizes were determined based on historical data from our laboratory using a power analysis to provide >80% power to discover 25% differences with p<0.05 between groups to require a minimum of 4 animals per group for all behavioral outcomes, and 2 animals per group for RNAscope analyses. All experiments were randomized by cage and performed by a blinded researcher. Researchers remained blinded throughout histological, biochemical, and behavioral assessments. Groups were unblinded at the end of each experiment before statistical analysis. All data are expressed as the mean ± SEM. Statistical analysis was performed using GraphPad Prism version 10.0.2 (GraphPad Software) or R, as described in Methods. Data were analyzed using two-tailed Student’s t tests, as indicated in the main text or figure captions, as appropriate. The “n” for each individual experiment is listed in the figure legends. A combination of male and female mice young or aged were used throughout the study. No data were excluded from analyses. In some cases, the different sections of the same DRG from the same mouse was used to detect multiple senescence markers for RNAscope analyses. To quantify RNA expression in DRG neurons, DAPI nuclear stain was used to determine glial-neuronal boundary to carefully associate RNAscope puncta with neuronal cell bodies. For mouse DRG, neurons expressing RNA were quantified per probe set as: IL6-positive >5 puncta; p16-positive >10puncta; p21-positive >20puncta; Trpv1-positive >20puncta. Human DRG neuron cut offs for positive counted cells were as follows: IL6 >10puncta, p16 >15puncta, p21 >20puncta, Trpv1 >20puncta. RNAscope quantification cell counts were done in a blinded fashion where experimenter was blinded to age/sex/timepoint per experiment. All counts were conducted assessing similar numbers of total DRG neurons per experiment.

### Data/Code Availability

Data that support the findings of this study are available from the corresponding author upon reasonable request. Publicly available code was implemented in Python for the UMAP (https://github.com/lmcinnes/umap) and HDBSCAN (https://github.com/scikit-learn-contrib/hdbscan) analyses. Figures in these analyses were generated using Matplotlib (https://matplotlib.org/) and Seaborn (https://seaborn.pydata.org/).

## ACKNOWLEDMENTS

We would like to thank Dr. Akila Ram, Andrea Cortez, and Laura Colman for technical help and discussion during the course of the project. Thank you to Dr. Amy Nippert and Dr. Heike Fuhrmann for helpful comments during manuscript preparation. We used BioRender software to prepare multiple schematics included in this paper.

## AUTHOR CONTRIBUTIONS

Conceptualization: LJD, VLT

Methodology: LJD, CLB, SFB, JAK, VLT

Formal analysis: LJD, CLB, SFB, OCG

Investigation: LJD, CLB, SFB, APL, LHH, CEJ

Writing—original draft: LJD, CLB, VLT

Writing—review & editing: LJD, CLB, SFB, APL, LHH, CEJ, OCG, LDL, JAK, VLT

Visualization: LJD, CLB, SFB, JAK, VLT

Supervision: LDL, JAK, VLT

Project administration: JAK, VLT

Funding acquisition: LJD, CLB, OCG, JAK, VLT

## SUPPLEMENTAL FIGURE LEGENDS

**Supplemental Figure 1. Confirmation of p16^INK4A^-specific RNA expression in the DRG.** RNAscope using RNA probes spanning exons encoding both p16^INK4A^ and p19^ARF^ protein (*Cdkn2a-tv1*). *Cdkn2a-tv2* RNA probe spans exons specific to p16^INK4A^ protein. Complete cellular co-localization of the *Cdkn2a* variants in mouse lumbar DRG sections confirming p16-specific expression in the mouse lumbar DRG. Scale bar of upper panels are 25µm. Scale bars of inset are 15µm.

**Supplemental Figure 2. Senolytic treatment does not alter normal sensory function.** The mechanical threshold of the contralateral (uninjured) hindlimb was measured after application of senolytic ABT263 or vehicle control in aged (A) and young (B) mice at Day19 post-treatment (n=21 aged vehicle-treated mice, n=24 aged ABT263-treated mice; n=9 young vehicle-treated mice, n=11 young ABT263-treated mice; two-tailed unpaired t-test, p= 0.8562 (aged mice), p=0.5858 (young mice)).

**Supplemental Figure 3. Increased percentage of human DRG neurons filled with lipofuscin with age.** A. Representative neurons in aged (65yr) human DRG with accumulated lipofuscin, a marker of senescence. Example of neuron either mostly filled (left arrow) or completely filled (right arrow). Scale bar is 10µm. B. Quantification of DRG neurons whose cell bodies were greater than 75% occluded by lipofuscin as a percentage of all DRG neurons in young and aged human DRG. Lipofuscin signal is defined by strong autofluorescence signal across all channels (488, 550, 647) which presents as a bright yellow/white signal in overlay.

**Supplemental Figure 4. SASP-expressing senescent neuron diameters in young and aged human DRG.** Cell diameters (µm) of human DRG neurons co-expressing either p21+IL6+, p16+IL6+, or p21+p16+IL6+, as a percent of total neurons counted in each population in young (A) and aged (B) DRG (Young (33yr) DRG: n=14 *p21*+*IL6*+ DRG neurons, n=78 *p16*+*IL6*+ DRG neurons, n=48 *p16*+*p21*+*IL6*+ DRG neurons; Aged (65yr) DRG: n=36 *p21*+*IL6*+ DRG neurons, n=48 *p16*+*IL6*+ DRG neurons, n=118 *p16*+*p21*+*IL6*+ DRG neurons).

