## Supplementary figures and images for "Aging and injury drive neuronal senescence in the dorsal root ganglia"

### Supplemental Figure 1

Supplemental Figure 1

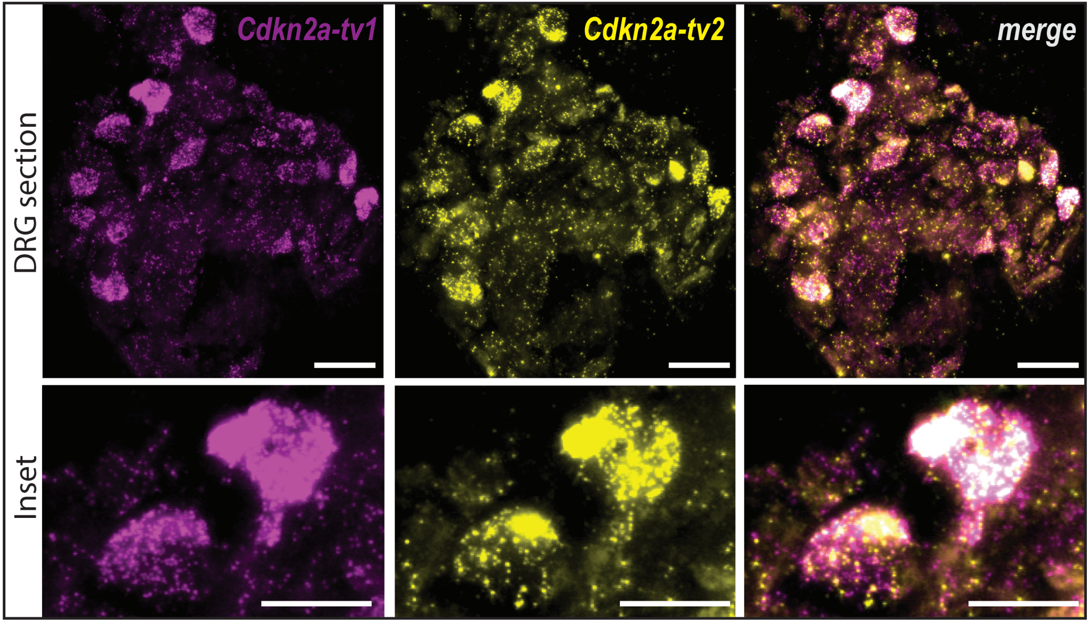

### Supplemental Figure 2

Supplemental Figure 2

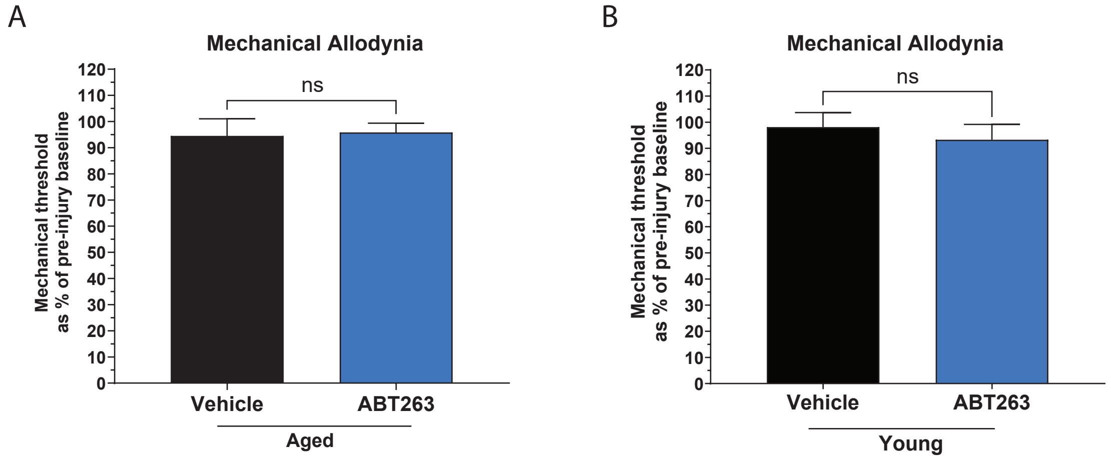

### Supplemental Figure 3

Supplemental Figure 3

A

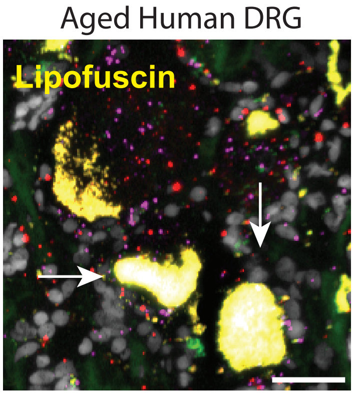


B

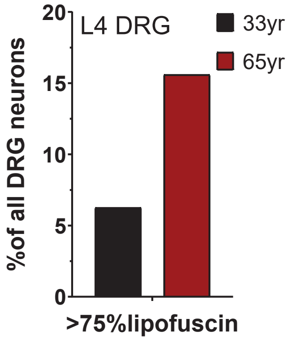

### Supplemental Figure 4

Supplemental Figure 4

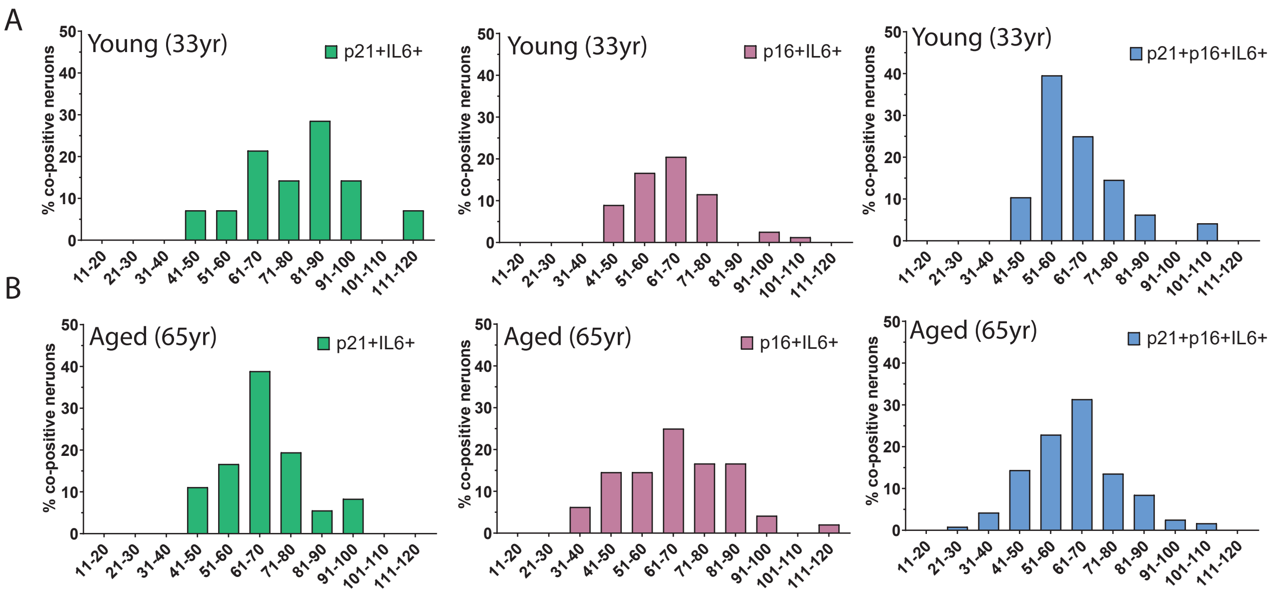
